# Cancer classification in the genomic era: five contemporary problems

**DOI:** 10.1101/023127

**Authors:** Qingxuan Song, Sofia D Merajver, Jun Z. Li

## Abstract

Classification is an everyday instinct as well as a full-fledged scientific discipline. Throughout the history of medicine, disease classification is central to how we develop knowledge, make diagnosis, and assign treatment. Here we discuss the classification of cancer, the process of categorizing cancer subtypes based on their observed clinical and biological features. Traditionally, cancer nomenclature is primarily based on organ location, e.g., “lung cancer” designates a tumor originating in lung structures. Within each organ-specific major type, finer subgroups can be defined based on patient age, cell type, histological grades, and sometimes molecular markers, e.g., hormonal receptor status in breast cancer, or microsatellite instability in colorectal cancer. In the past 15+ years, high-throughput technologies have generated rich new data regarding somatic variations in DNA, RNA, protein, or epigenomic features for many cancers. These data, collected for increasingly large tumor collections, have provided not only new insights into the biological diversity of human cancers, but also exciting opportunities to discover previously unrecognized cancer subtypes. Meanwhile, the unprecedented volume and complexity of these data pose significant challenges for biostatisticians, cancer biologists, and clinicians alike. Here we review five related issues that represent contemporary problems in cancer taxonomy and interpretation. 1. How many cancer subtypes are there? 2. How can we evaluate the robustness of a new classification system? 3. How are classification systems affected by intratumor heterogeneity and tumor evolution? 4. How should we interpret cancer subtypes? 5. Can multiple classification systems coexist? While related issues have existed for a long time, we will focus on those aspects that have been magnified by the recent influx of complex multi-omics data. Ongoing exploration of these problems is essential for data-driven refinement of cancer classification and the successful application of these concepts in precision medicine.

## Introduction

Classification and labeling represent the most intuitive forms of learning. According to the Confucius, “If the name is not right then speech will not be in order, and if speech is not in order then nothing will be accomplished”. Instances of classification can be found in every aspect of life: the Linnaean system in biology is a seven-level framework for cataloging living organisms; E-commerce companies must organize their holdings in an easy-to-search system. Whether the goal is to recommend movies, screen job candidates, or rank colleges, a system of summarizing and dividing is involved, and uncertainties inherent to this task are common. In a quote attributed to Albert Einstein [1], classification is “an attempt to make the chaotic diversity of our sense experience correspond to a logically uniform system of thought”. The tension between “chaotic diversity” on one hand and a “logically uniform system” on the other makes classification a long-standing subject in scientific research. In practice, it was regarded more often as an art than a science, as there is no axiomatic classification theory that applies to all problems. This review will discuss the topic of cancer classification. Instead of offering a systematic review of specific cancers we will focus on the implicit assumptions and common caveats of classification, especially how it has been confronted with new challenges in the genomic era. We organize the many strands of this topic under the heading of Five Contemporary Problems, although in many ways they are new, more acute forms of old problems. A recurring theme of our discussion is that the merit of a classification system depends on what it will be used for. Often, a new map is drawn, but its intended application is unclear. Only by spelling out the purpose of a proposed system can we evaluate its merit according to what it aims to achieve. The potential measures of merit for classification systems include: statistical robustness, connections with existing standards, prognostic value [2], insights into biological mechanisms, and predictive power of treatment outcomes [3]. As we discuss below, sometimes these merits cannot be achieved all at once; and multiple classification systems can be simultaneously “correct” depending on which purpose they are designed to serve.

A few notes of clarification before we begin. In the field of statistical learning the term “classification” refers to *supervised sample assignment,* using features identified in a training set containing samples with known class labels. Here we adopt its other, more commonly understood meaning, referring to *ab initio* pattern recognition, also known as *unsupervised class discovery.* We will use the terms (sub)class and (sub)type interchangeably and will only discuss sample classification, not gene clustering. We sometimes use “genomics” to refer to all high-throughput-omics data. We will focus on subtype discovery within a major organ-specific cancer type, not between the major types.

## 1. How many cancer types are there?

The short answer is about 200: the National Cancer Institute keeps an A-Z list of nearly 200 cancer types [4], organized by organ location, although with some exceptions, such as “HIV-related” or “unknown primary”. This organ-centric system is further stratified, most often by the known cell type of origin within the organ, e.g., “astrocytomas”, as a subtype of brain tumors [5], or by patient age, e.g., “childhood leukemia”. If each of these major types has four subtypes, there will be 800 subtypes of cancer. As stated above we will focus on how to identify subtypes within a broadly recognized organ-specific entity.

### “Endless forms most beautiful”

While the organ-derived naming system has been used for over a century by doctors, patients and cancer registries, there has always been the drive to recognize finer subtypes; and this trend has accelerated dramatically with genomic data. For example, in the clinic, primary breast cancers have traditionally been classified by their expression status of hormone receptors into three subtypes: estrogen receptor (ER)-positive or progesterone receptor (PR)-positive, human epidermal growth factor receptor 2 (HER2)-positive, and triple negative [6]. The result of the immunohistochemical (IHC) assays of these receptors can thus guide the selection of patients for different hormonal therapies or targeted therapies [7, 8]. The arrival of microarray-based gene expression data led to the division of breast cancers into five intrinsic molecular subtypes: luminal A, luminal B, HER2-overexpression, normal-like, and basal-like, and they showed notable differences in clinical outcome [9, 10]. More recently, analyses of copy number and gene expression data for several thousand malignant breast tumor samples revealed 10 molecular subtypes [11, 12]. Meanwhile, the triple negative subtype was further divided [13, 14]; and with a sample size of >10,000 and additional basal markers Blow et al. [15] identified six IHC subtypes for breast cancer by splitting the basal group.

At least three factors contribute to the continued fracturing of cancer subtypes. First, increasingly larger sample cohorts are being used—they contain not only more phenotypic heterogeneity but also greater power to define and replicate rare subtypes. Second, simultaneous collection of diverse-omics data types, for aberrations in DNA, gene expression, epigenetic features, etc, tend to recognize additional “structures” in a given tumor series [16-18]. Third, intrinsic gene-gene correlations in the data, often unrelated to the disease itself, may create illusions of stable clusters with the use of some methods [19] even in cases where the structure is weak.

An ever finer classification system has many potential benefits. It is needed to capture the full spectrum of biological diversity—the “endless forms” that Darwin spoke of. It could lead to a better recognition of patient-specific disease mechanisms, and importantly, could suggest treatment options that are more accurately matched to the patient’s tumor [2, 3]. Precision medicine, at its very foundation, relies on valid and constantly optimized disease classification that reflects the underlying mechanisms. However, a finegrained classification system also has many potential drawbacks. The newly proposed splits may not be technically robust (see #2 below). Even when the finer categories are robustly supported by statistical significance and by replication, they may still lack a clear biological meaning, or have little impact on treatment options (#3-4 below) if it turns out that some subtypes share the same clinical endpoint, or if treatment options are limited.

## 2. Is the classification system robust?

Here we delve into the issue of robustness: how can we tell if a classification scheme is more reliable than alternative systems? The value of any classification depends critically on its robustness. Without evaluating robustness we face profound problems in downstream analyses, including integration across studies and clinical translation. For example, the reported number of subtypes for a certain cancer could differ between two studies, simply because one or neither study had strong evidence to formally distinguish K clusters from (K-1) or (K+1). Similarly, if the reported optimal K vary among DNA, mRNA, and methylation data, the discrepancy could either reflect a real biological distinction, or be explained by trivial methodological differences or by the mere absence of a strong cluster signal.

### Is there a p value?

In epidemiological or genetic association studies, evidence of credible association is measured by effect size and statistical significance, the latter being expressed by a p-value and a hypothesis-testing procedure used to calculate it. For example, a DNA variant’s additive effect on a continuous trait can be evaluated by linear regression. However, the task of classification cannot be easily cast into a hypothesis-testing framework: when declaring K clusters for a sample, is the null hypothesis “no cluster” or “K-1 and K+1 clusters”? While the confidence of *class assignment* can be assessed by cross-validation in test samples for which the class labels are already known, there is no well-established statistics to compare the performance of *class discovery.* In theory, one can quantify the degree of clustering by indices such as compactness, connectedness, separation [20], or silhouette width [21], however there is no universally accepted quantitative measure – a p value-like index - to report how likely the observed clusters could arise merely due to naturally occurring data “structure”. Two types of structure are frequently encountered in high-dimensional molecular profiling data: that due to separations between groups, i.e., stratification, and that due to locally tight clusters, i.e., cryptic relatedness. These terms are borrowed from human population genetics studies, where both types of structure ultimately came from shared ancestry of sampled individuals at different time depths. Their impact on association tests can be monitored and corrected by well-established procedures [22, 23]. However, for gene expression or other functional genomics data (such as proteomic, metabolomic, epigenomic data), the information used in classification is sample-sample similarity in high-dimensional feature space, and the basis of co-ancestry is lacking, at least not self-evident. Indeed, how to evaluate competing algorithms or alternative results is an active topic of research [24]. Many groups have studied the issue of cluster validation and have proposed the use of either internal or external standards [25-27]. More often however, there is no real dataset that can serve as a reliable external standard. Our recent analyses have shown that even the datasets that are said to contain well separated clusters can have an uncertain number of clusters (i.e., the true K), thus making it difficult to use them as benchmarks for comparing class discovery methods [19]. As sample size increases, the number of clusters will increase and exhibit increasing apparent stability [19, 28]. Given this, we recommend the use of simulations to create fully transparent standard data, in which the number of clusters (K) and the degree of separation are known, and the gene-gene correlations can be introduced based on empirical data. Using such simulated data as a benchmark we can systematically compare different clustering methods for their performance, especially their rate of over- and under-calling K over data that span a wide range of known K values and pre-specified degrees of cluster separation.

Quantitative reporting of the robustness of clustering results is often lacking in publications that propose new classification systems. Sometimes the data structure was *illustrated* by pre-selecting the best discriminating genes and showing how they could visually separate the reported clusters crisply.

Although this form of presentation is well suited for annotation - showing which genes appeared in which group - it is not appropriate as a demonstration of cluster strength, because with many more genes than samples (i.e., the p >> n situation) seemingly informative discriminators can always be found for any random partition, even for samples without clear groupings. When classification strength is not properly assessed, visual display of clusters using the best genes can inadvertently turn into an exaggerated inference, even if subsequent interpretations seem appealing [19].

## 3. Can classification capture intratumor heterogeneity and evolutionary progression?

Every living cancer inevitably changes its character in time and every solid tumor is spatially heterogeneous, yet most samples used in research so far are bulk tissue blocks collected as a single time point. Thus most of today’s cancer genomics data, by the very nature of sampling, provide a one-time view of a mixed pool of changeable cells. Standard cancer classifications are aimed at capturing *intertumor* heterogeneity, while treating each tumor as homogeneous and unvarying. For a given cohort of patients, classifying their bulk tumor data is an attempt to find natural groupings among many mixed cell populations, while ignoring the within-population diversity and its variation in space and time. Not surprisingly, the cancer subtypes reported are often driven by, and therefore reflect, intratumor heterogeneity and tumor evolution.

### Spatial heterogeneity

With bulk-tissue data, spatial heterogeneity is an unobserved property. If the sample consists of a limited number of clonal populations, the number and molecular features of these component populations can be potentially “deconvolved” computationally. As sequencing costs drop, spatial heterogeneity can be analyzed with increasing resolution: first by multi-region analysis of smaller sectors of the same tumor [29-31], ultimately by DNA or RNA sequencing of single cells [32, 33]. In multi-region analyses, a typical assumption is that individual regions are clonally “pure”, and the results can be presented as such. However, it remains the rule rather than the exception that each region still contains a mixture of many cell types, albeit with presumed lower heterogeneity compared to the entire tumor [34]. Single-cell analysis provides the ultimate solution, as it describes the smallest unit of cancer heterogeneity and provides truly clonal data for use in classification. Single-cell studies have identified many more cell types than previously seen with bulk tissue analysis. For example, a single-cell RNAseq study of cortical tissues [35] has found nine major classes and 47 “molecularly distinct” subclasses of brain cells, significantly expanding the known cellular repertoire of the mammalian cortex. Due to the higher cost of single-cell analysis, bulk tissue samples will remain the predominant source of cancer genomics data for the foreseeable future, both in basic research and in real-time clinical testing. By sampling smaller and smaller “core” regions the spatial heterogeneity can be reduced, but it cannot be fully removed. In this regard the traditional, *hard* classification into disjoint categories is a poor fit for admixed samples, as they contain cancer cells carrying somatic mutations or aneuploid segments as well as surrounding normal-like cells that are euploid and carrying only germline mutations. Partial membership modeling has been proposed to address this scenario [36, 37], reminiscent of similar methods for ancestry inference using genotype data for individuals with mixed ancestry [38, 39]. Alternatively, phylogeny-based methods can be applied to explicitly account for the polyclonal nature of each regional sample in a multi-region dataset [40, 41].

### Cancer life history and impact on classification

Classification of cancers can also be viewed as a task of cataloging evolutionary trajectories of complex genomes [42, 43]. Each cancer genome carries many variations, with a distribution of fitness effects as defined in a specific but changeable environment. Each tumor thus undergoes its own Darwinian evolution, with many intricate details that make it distinct from all other tumors [44]. However our effort to classify them is predicated on the notion that convergent evolution does happen, such that a limited number of evolutionary paths are traveled repeatedly by tumors from different patients, leading to recognizable major hallmarks recurring in different tumors. The discovery of N subtypes of breast cancer, for example, detects N destinations of convergent evolution in this cancer type. But N needs not be a fixed parameter. For example, if the tissue has K cell types that could eventually turn to cancer cells, the pre-cancerous cells have K starting points from which to explore the initial evolutionary paths, and multiple paths may merge or split during the life history of a cancer before congregating to one of N major types at the time of sampling. Metastasis and treatment response would further extend, diversify, or reshuffle the evolutionary trajectories [45, 46].

Current cancer classification systems can only reflect, but not accurately portray, this complex succession of events, as different tumors may be sampled at different stages along their own life histories. In a typical lifespan of a malignant tumor, some cells start acquiring oncogenic potential, showing increased proliferation, escaping apoptosis and immunosuppression, competing successfully for resources with other cells in the tissue niche, harnessing the right combination of driver mutations with enough fitness gain to overcome the fitness burden incurred by the larger number of passenger mutations [for reviews of common hallmarks and pathways for cancer, see 47, 48, 49]. To become more “successful” the cancer needs to grow sufficiently large and clonally diverse before tissue turnover, and sometimes, to acquire migration and invasion properties that are essential for metastasis. It may also carry treatment-resistance subclones that can thrive after therapy. Viewed in this light, cancers are chronic diseases with changeable character, punctuated by rare episodes of acceleration, essentially slow-evolving populations that occasionally rush to a new, more successful endpoint. A given sample collection may have captured the tumors at different “stations” of their life history. For example, the mesenchymal subtype of glioblastoma multiforme (GBM) possesses the signature of macrophages/microglial infiltration [50, 51], and has a greater degree of mixing of aneuploid and euploid cells. More recently, Ozawa et al. reported that most non-GCIMP mesenchymal GBMs may evolve from another known subtype, a proneural-like precursor of GBM [52].

## 4. Caveats in interpreting cancer subtypes.

In most cases, the task after class discovery is to explain the meaning of the found classes. Typical actions include: describe the involvement of known cancer-related genes as a way to report specific signaling pathways active in different subtypes; map the new class nomenclature to those of the previously established systems; assess differences in clinical outcome, e.g., survival time or treatment responsiveness; dissect tissue-specific signatures or enrichment of previously curated gene clusters. In the following sections we discuss three caveats in such interpretations. Both marker selection and analysis method have a strong impact on classification results and how the resulting classes can be explained. Further, the third factor affecting interpretation is the strength of clustering inherent in the dataset, which is not known until a specific collection of tumors has been assembled and analyzed.

### Feature selection bias

We illustrate the importance of feature selection by first drawing analogies with population genetics studies, where DNA variation data can be used to (1) infer historical demography or (2) detect natural selection. These two tasks are related, but are also inherently different, even conflicting with each other, and relying on different features. Demographic inference tries to describe historical changes of population size and distribution, including migration, self-isolation, expansion, and admixture. The best genetic features for this task are “neutral” variations, those not under natural selection, or more accurately, those not likely to affect the Darwinian fitness of the individuals. Typical examples of such neutral markers are those found in intergenic regions of the genome. In contrast, the second task, inference of natural selection, relies on markers with a fitness effect and they can be detected after specifying a null model of genetic drift, because demographic forces could have generated DNA variation patterns similar to those due to natural selection.

In cancer evolution, the concept of *driver* and *passenger* mutations are almost exact parallels of the *adaptive (positively selected)* and *neutral* variants recognized in population genetics [49, 53]. However, we rarely ask whether the emphasis of cancer classification is for understanding the tumors’ past demography, or their future evolution. The story of the cancer’s past is shaped by the drivers but may be best recorded by the passengers. And the drivers in the past may not experience the forces of selection acting in the future. The type of features selected for use in classification will therefore directly affect the downstream interpretation. For example, if the analysis of gene expression data pre-selects transcripts that correlate with survival time, these markers, by being most informative for future outcome, are likely to reveal tumor subtypes that differ in outcome, and this in turn will lead to the interpretation that the subtypes thus discovered have a stronger prognostic/predictive value than classes discovered using other features. Alternatively, if the selected transcripts correspond to cell type or pathway-specific genes, the resulting classes will likely exhibit differential loading of pathway or source cell signals, and make it easy to map the discovered classes to biological mechanisms. Even the presumably “unbiased” choice of selecting the most variable genes may have inadvertently loaded the interpretation towards the most dynamic or most strongly co-expressed pathways, such as those for stress response or immune cell infiltration.

### Hidden assumptions in choosing a clustering method

Unsupervised clustering is the basic tool for *ab initio* class discovery. The term “unsupervised” refers to statistical inference of data structure without relying on existing knowledge of sample labels. However, “unsupervised” does not mean assumption-free. Some important assumptions have always been made, sometimes unknowingly, in marker selection, data processing (such as how to treat outliers and how to deal with batch effects), and the choice of the clustering method. This methodological choice is already based on an implied data-generating model, i.e., what type of biological heterogeneity could have produced the observed data structure. If we assume that the objects - different tumor samples - arise from distinct, non-overlapping causes, methods for finding disjoint taxonomy, such as k-means clustering, are appropriate. If, alternatively, we see each tumor sample as a mixed population of cells comprising a small number of canonical clones, a mixed membership model is more powerful [36, 37, 54], as has been routinely applied in studies of human population diversity involving individuals of mixed ancestry [38]. The number of co-existing clones and the rate of clonal replacement depend on population size, mutation rate, and the distribution of fitness effects of the new mutations. If the task is to classify cells in a single evolving population with major branches, it would be best to capture the lineage relationship by using hierarchical clustering - akin to the coalescence analysis in classic population genetics [55] - aiming to identify hierarchical classes to reflect their shared ancestry as the historical truth. When using simulated data to compare methods such as k-means clustering or hierarchical clustering, the data-generating model will usually dictate the winner: the method that matches the model will fit the simulated data best.

### Inherent intensity of data structure

Perhaps it is worth reiterating that when the clustering signal is strong, most methods will perform well and they will be in good agreement. Conversely, when the data structure is subtle, slight differences in sampling, data processing, feature selection, or the choice of method will yield highly discordant results, and it is difficult to tell which method is better. A recent example of strong clustering signal is from a joint analysis of 3,527 specimens of 12 cancer types by the Cancer Genome Atlas (TCGA) Research Network [17]. Most identified classes follow the cancer’s known organ of origin, undoubtedly due to the distinct cell types found in different tissues. This result is as expected, because it affirms the earliest insight that cancer cells are transformed normal cells rather than entirely “foreign” cells (as is the case in bacterial infection). As cells are “canalized” during differentiation, a tumor occurs in one of the Waddington’s valleys and bears its local hallmark. Speaking in more modern terms, the malignant transformation, although a radical step in the adaptive evolution of the cancer cells, usually could not have erased their tissue-of-origin signature inherited from prior differentiation. This identity, perhaps coded in the epigenomes of the fate-committed cells, remain the most noticeable molecular character in matured organs despite subsequent oncogenesis. Meanwhile, this study also revealed remarkable examples of cross-tissue subtypes [17], e.g., squamous-like lung, head and neck, and bladder cancers that are more similar to each other than other cancers from the same organ, and they appear to have overcome the ontological divergence of the source tissues. Is this because convergent evolution—newly acquired oncogenic characters over-writing history, or shared lineage—the same group of differentiated cells “seeded” into two different body sites? Either scenario would be immensely interesting. In principle, it is entirely possible that the same pathways are activated in different cell types as the oncogenic driver and manifest as the same subtype. Such cross-tissue classification is at the forefront of pan-cancer analysis today, made possible by the availability of multi-cancer datasets collected under uniform technical conditions. With such data we are entering the best time to study population genetics of somatic cells. This discipline will stimulate new theoretic work on the evolution of non-recombining populations, produce patterns that can be contrasted with experimental evolution of microbial systems, and will provide increasingly strong knowledge support for the practice of precision oncology.

## 5. Can multiple classification schemes coexist?

Traditional classification methods rely on organ type, appearance, and histological markers. Multiple systems, such as tumor grade and stage, have coexisted for decades. In recent years, the arrival of high-throughput molecular studies has rapidly increased the number of competing systems. For example, the analyses of human breast tumors by the TCGA [56] produced multiple answers to the “how many subtypes” question for the same sample series. It found that breast tumors’ gene expression data supported 13 classes with the use of a consensus cluster-based method, 12 classes with a second method, and five classes with the semi-supervised PAM50 method [57]. The concordance rate among the three results was modest, as the best-matched classes between any two methods only account for 50-60% of the samples (our unpublished observation). Further, the study found seven breast tumor subtypes from microRNA data, five subtypes from methylation data, and five subtypes from copy number alterations, again with poor to modest agreement (per our analysis). While the classification solutions were described as “correlated” across data types, their differences impacted a large fraction of the samples, making it difficult to find a straight answer to the simple question: how many subtypes are there for breast cancer? To integrate such complex data requires quantitative assessment of clustering strength within each data type, and a system to truly integrate the solutions rather than recounting them side-by-side. Multiple methods have been developed to meet this growing need [58-61].

When the same data type lead to two or more different classifications by the use of different methods, it becomes a methodological imperative to arrive at a consensus: there is no sound reason for multiple “truths” within the same raw data. However, across different genomic data types it is less clear that there must be a single unifying classification. In the example of TCGA breast tumor study mentioned above, it may not be possible, or necessary, to forge a single classification by reconciling the observed groupings across all omics layers. If we assume that biological information flows from DNA to epigenetic marks, then RNA, and then to proteins, if the cells commit to its differentiated fate primarily using epigenetic codes yet react to short-term needs by gene expression adaptation, further, if the cells interact with their microenvironment chiefly through variations of metabolites, it is conceivable that different levels of biology coalesce to different grouping patterns, and the same sample could truly belong to different groups depending on the level of inquiry. We think it remains an open question whether different layers of genomic data signify different archetypes of cellular states and could lead to different but equally valid classification systems. To test such a multi-layered classification would require very large datasets that can validate the class-to-class mapping across layers.

## Conclusions

Cancer classification is both a scientific technique and a living art, to be performed for each dataset with individualized care. Classification results are widely used, and form the foundational knowledge for both basic and translational oncology. In this review we outlined five contemporary challenges at the interface of computational data mining, biological understanding, and clinical utility. To classify late-stage tumors is to engage in probabilistic cataloging and reverse story-telling, not unlike other observational sciences such as archeology or anthropology. The arrival of genomic data has dramatically increased the power to peer into the past, but even now, amidst the excitement of many new opportunities, it is useful to keep in mind that sometimes the sample series at hand may not be sufficient to support the full ambition of finegrained classification or to trace the entire evolutionary trajectories. Whether intratumor heterogeneity is so strong as to destabilize intertumor classification, and whether a newly observed cluster represents a robust, recurrent, and meaningful subtype, ultimately need to be settled by empirical data for each major tumor type. The usefulness of classification ultimately rests on increased prognostic power or more precisely targeted therapies. Assessing and communicating the strength of data, in quantitative terms whenever possible, is essential for the long-term management of predictive uncertainty, and for the successful application of genomics in patient care.

## Authors’ contributions

QS and JZL conceived the article and took the lead in writing the initial draft. All authors contributed to the research discussed in the article.

## Acknowledgements

QS is supported by funding from the Joint Institute for Translational and Clinical Research of the University of Michigan Health System and Peking University Health Sciences Center. SDM is supported by the Breast Cancer Research Foundation and the Metavivor Foundation. JZL is supported by general funds of the Department of Human Genetics and the Department of Computational Medicine and Bioinformatics of the University of Michigan. We thank two anonymous reviewers for their insightful comments.

